# Quantitative Substrate Kinetics Screening: One-Pot LC/MS Based Approach to Map Protease Specificity

**DOI:** 10.1101/2025.10.06.680748

**Authors:** Aaron E. Robinson, Garret D. Bland, Jessica Chu, James Ireland, Lilian Dao, Trinh Pham, Ming Dong, Vadim Demichev, Eric Johansen, Volker Schellenberger

## Abstract

Proteases play critical roles in many biological processes and diseases. Proteases tend to have limited and overlapping specificities, which hinders our understanding of their roles in physiology. We designed a naïve protease substrate library with the aim of fully utilizing modern mass spectrometry proteomics tools to quantitatively map protease substrate specificity. A library of approximately 200,000 peptides was efficiently produced in *E. coli* with sufficient library diversity to identify cleavage efficiency of thousands of substrates for most proteases in a single assay. Each experiment results in ample substrate kinetics data to build a positional specificity matrix that quantitatively maps protease specificity. We apply this method to two different proteases, GluC commonly used as a proteomics tool, and Fibroblast Activation Protein Alpha that is overexpressed in activated fibroblasts of epithelial cancers and fibrotic diseases. This research quantitatively maps substrate specificity for GluC and FAP in a single assay and can be applied to map substrate specificity for most proteases.

**Graphical Abstract:** 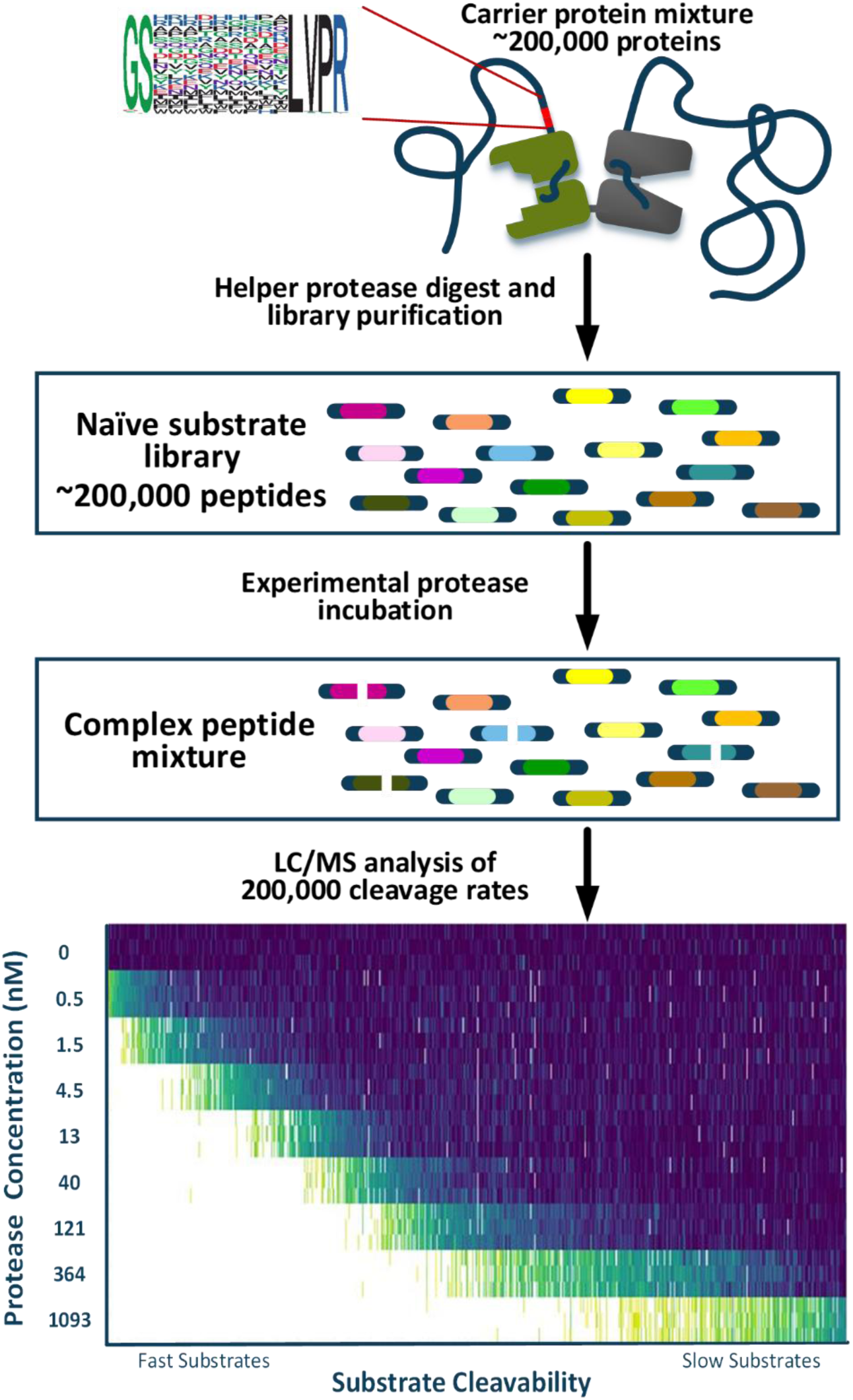

## Introduction

Proteases play an essential role in the human body and regulate key biological processes including cancer, inflammation, apoptosis, and protein degradation. ^1–4^ There are at least 553 proteases and homologues in the human degradome, representing 2% of the genome and they are grouped into five major families; aspartate, threonine, cystine, serine, and metalloproteases, each with peptidase activity evolved to fit its unique function.^1,5^ The intertwined web of proteases and their overlapping substrate specificities is complex and despite significant work in the field has yet to be detangled.^6–8^ In important step towards that goal of understanding the role of proteases is the precise mapping of their specificity. Prior efforts to map protease specificity have included techniques such as positional scanning peptide libraries,^9^ multiplexed peptide profiling,^10,11^ proteome-based discovery screens,^12^ phage-^7,13,14^ and RNA-display,^15^ each approach with its own set of pros and cons.

These innovations increased the number of substrates measurable within an experiment. Phage- and RNA-display can examine a large search space and are frequently used to assay protease substrate specificity, ^7,13–15^ but yield semi-quantitative results. Display approaches result in a holistic map of protease specificity, providing enriched substrate sequence motifs and require follow-up experiments with synthetic peptides for each protease to determine cleavage location and cleavage kinetics.^5,12^ Liquid Chromatography Mass Spectrometry (LC/MS) based approaches can directly measure substrate kinetics but tend to be limited by the number of substrates that can be investigated simultaneously and consequently library diversity. Positional scanning peptide libraries must be customized to individual proteases or groups of proteases with known starting substrates.^9,11^ Proteome based discovery screens^12^ and existing multi-substrate profiling approaches^10,11^ lack amino acid and sequence diversity necessary to quantitatively map a protease. In addition, libraries derived from cellular proteomes rely on peptides found in nature with unalterable length and hydrophobicity, inherently have a large dynamic range,^16^ and cannot distinguish leucine/isoleucine isomers, all limiting factors for assessment by LC/MS.

Here we describe a novel peptide library that was designed for optimal quantitation by LC/MS. Exposure of this library to proteases results in kinetic curves for thousands of substrates in a single assay. The resulting data is sufficient to train a positional specificity matrix (PSM) that quantitatively models protease specificity and can predict cleavage speed and location of most peptides for their respective proteases. This method brings together the precise quantitation of peptide degradation kinetics found in synthetic peptide library LC/MS screens^9,11^ and the broad substrate diversity found in phage- and RNA-display naïve libraries ^7,13–15^ to quantitatively map substrate specificity of a protease in a single assay.

We applied this approach to two proteases, GluC and Fibroblast Activation Protein Alpha (FAP). GluC is a serine protease isolated from Staphylococcus aureus and that is commonly used as a proteomics tool cleaving peptide bonds C-terminal to glutamic acid and aspartic acid residues.^17,18^ FAP is a human protease known to be overexpressed in activated fibroblasts of epithelial cancers and fibrotic disease tissues. Despite the biological relevance of FAP and its physiological dual function as both an endopeptidase and dipeptidyl peptidase, its substrate repertoire is not fully understood with only 10 substrates reported in the MEROPS database of protease substrates.^19–21^

The method established here quantitatively maps substrate specificity for a protease in a single LC/MS experiment without any prior knowledge of that protease’s specificity with the quantitative precision needed to identify subtle differences in structurally similar proteases.^6,22,23^

## Results

### Designing a Naïve Peptide Substrate Library

Designing a universal peptide substrate library optimized for LC/MS with the ability to determine substrate specificity for a wide range of proteases was the first and arguably most important step. We chose to use a naïve library similar to phage- and RNA-display^7,13,14^ and deliberately optimized peptide sequences for diaPASEF acquisition.^24^ Helper proteases Thrombin and Factor Xa were used for their narrow substrate specificity to cleave the substrate library from its carrier protein, resulting in C-terminal arginine containing peptides. Cystine residues, substrates with 4 or more of the same amino acids in a row, isomers, and substrates where leucine and isoleucine would be undifferentiable in LC/MS experiments were excluded from the design. Prosit^25^ and DeepCCS^26^ were used to predict retention time and ion mobility distribution for of all possible substrates. 90,000 substrates were chosen for the Factor Xa and Thrombin sub-libraries with a uniform retention time distribution, −10 to 150 in the cIRT^27^ normalized retention time space, and a uniform ion mobility distribution while maintaining sufficient peptide diversity when combined (Figure S1A-D, Table S1).

### Generation, Purification, and Characterization of Substrate Library

Library sequences flanked by canonical Thrombin or Factor Xa helper protease sites (LVPR-GS and IEGR-G, respectively) were cloned as a pool into the unstructured XTEN polypeptide region to the N terminus of an EGFR Bi-specific T-cell engager,^26^ that served as a carrier protein (Figure 1). After cloning, confirmatory NGS was performed resulting in the identification of unique 232,199 substrate sequences with more than three reads, including 99.7% of the designed sequences. (Figure S2A-B). The mixture was fermented in *E. coli* and purified as previously described^28^ and digested into a peptide library using helper proteases Thrombin or Factor Xa. The complex product mixture was further processed to remove helper proteases and domains of the carrier protein (Figure S3). The two sub-libraries were combined resulting in a high purity naïve substrate peptide library used for the subsequent assays.

**Figure 1:**
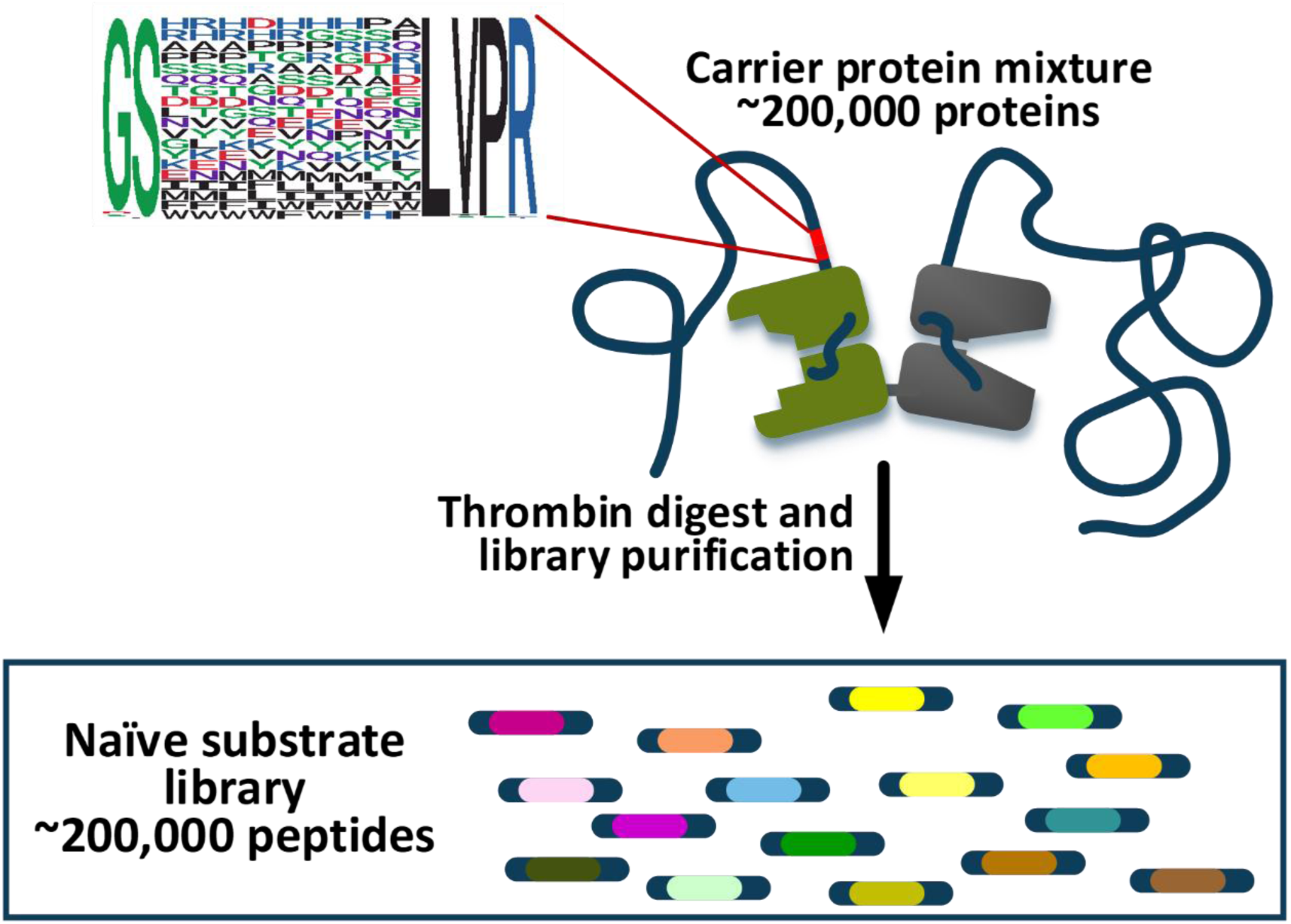
Naïve substrate library was expressed in *E. coli* and purified as a protein mixture. Protein mixture was exposed to a helper protease Thrombin or Factor Xa to cleave the library region from the carrier protein. The substrate library was purified away from the carrier protein and helper protease was removed resulting in a naïve substrate library of ∼200,000 peptides.

Substrate library composition was assessed without the addition of a protease to characterize the library and optimize LC/MS methods. Since the naïve substrate library was purposefully designed, very little optimization was required, and a uniform retention time and ion mobility distribution was observed with a standard 100min LC gradient and diaPASEF method (Figure S4A-B). After library-free analysis of triplicate injections with DIA-NN 2.0,^29^ 92,595 substrates were identified. Of those, 73,479 substrates were quantified with less than 20% CV (Table S2).

### Mapping GluC’s Substrate Specificity in a Single Assay

To assess naïve substrate library performance, we performed a cleavage experiment with GluC by incubating the substrate library without protease and with 8 incrementally increasing concentrations of GluC for a fixed incubation time (Figure 2A) and acquired triplicate diaPASEF injections for a total of 27 acquisitions.

**Figure 2:**
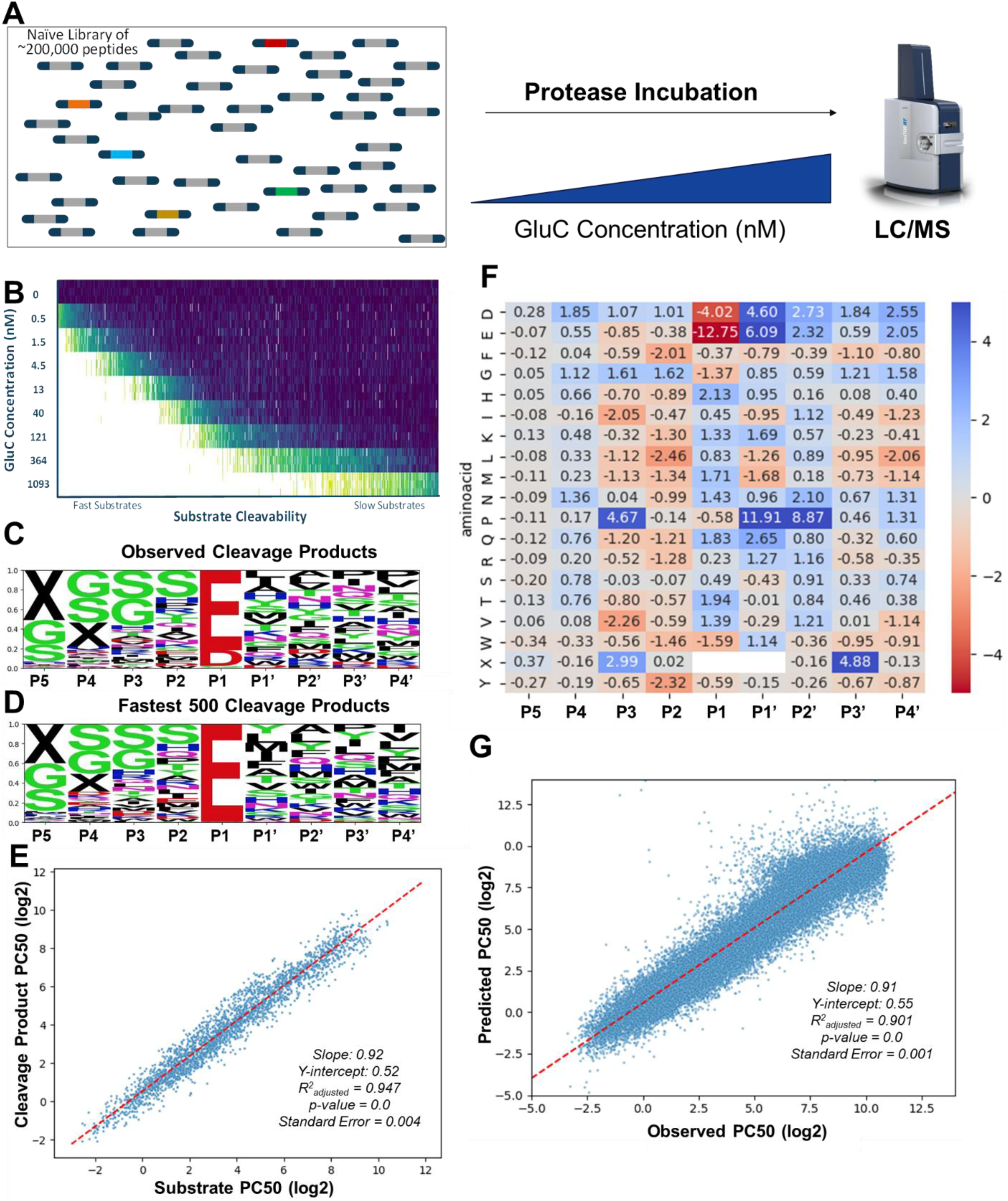
**(A)** Experimental workflow schematic where we expose our naïve substrate library to a GluC dilution curve before running diaPASEF LC/MS acquisitions **(B)** Heatmap of high quality cleaved substrates in GluC substrate kinetics experiment showing a gradation of very fast substrates to relatively slow substrates cleaved by GluC **(C)** SeqLogo of 2,850 peptides cleaved by GluC showing an enrichment of glutamic acid and aspartic acid in P1 **(D)** SeqLogo of the fastest 500 peptides cleaved by GluC showing an enrichment of glutamic in P1 and hydrophobic residues in P1’ **(E)** Scatterplot of 2,850 substrates where we could detect high quality products (Y Axis) and high quality cleaved substrates (X Axis) showing the correlation of PC50 measurements of each. 93 pairs with absolute difference > 2 were considered outliers **(F)** Positional Scanning Matrix (PSM) of GluC substrate kinetics experiment characterizing GluC’s substrate specificity. All amino acids are fixed to alanine; a negative number represents the log2 PC50 speed an amino acid substitution will speed up a substrate and a positive number will slow down the substrate in respect to alanine. X is indicative of a lack of a residue **(G)** Scatterplot showing PSM predicted PC50’s (Y-Axis) vs Observed PC50’s (X-Axis) for the GluC substrate kinetics experiment with the naïve substrate library. Predicted PC50’s correlated with observed PC50’s with a R^2^=0.901 and slope = 0.91.

Recent improvements in the DIA-NN software suite, including QuantUMS^30^ in DIA-NN 1.9.1 and custom decoy implementation for peptide libraries with fixed N- or C-terminal amino acids in DIA-NN 2.0, have resulted in significant improvements in data quality for this dataset (Supplementary Information, Table 1, Figure S5A-F). This effect is magnified further after subsequent secondary data analysis and DIA-NN 2.0 resulted in significantly more substrates cleaved faster by GluC (Figure S5C, Table 1). Therefore, we chose to use DIA-NN 2.0 for primary data analysis of these complex datasets moving forward.

**Table 1:** DIA-NN 1.8.1, 1.9.1, and 2.0 were used to process the GluC substrate kinetics experiment. After DIA-NN processing, each went through the same secondary analysis pipeline and substrates identified are split into 5 categories, High Quality Cleaved (HQ-CLV), Cleaved (CLV), High Quality Not-Cleaved (HQ-NOC), Not-Cleaved (NOC), and Unknown. HQ-CLV substrates fit the first order rate equation with a significant p-value (>0.05), are observed in triplicate no protease acquisitions, must have a minimum of 6 datapoints, and be observed in at least 3 unique protease concentrations. CLV Substrates fit the first order rate equation with a non-significant p-value (<0.05), are observed in triplicate no protease acquisitions, must have a minimum of 6 datapoints, and be observed in at least 3 unique protease concentrations. HQ-NOC substrates that fit a straight line with p-value > 0.05 and no missing datapoints in triplicate LCMS acquisitions at each protease concentration. NOC substrates fit a straight line with p-value < 0.05 and no missing datapoints in triplicate LCMS acquisitions at each protease concentration. Unknown substrates don’t fit either the first order rate equation or a straight line. High Quality products (HQ-products) were obtained by a second DIA-NN search looking for all full-length substrates and for all possible products coming from HQ-CLV substrates and fitting all products to a modified 2-parameter fit of the inverse of the first order rate equation with a p-value < 0.005 which match to a unique HQ-CLV substrate.

| GluC Substrate Kinetics Assay: Optimizing Data Quality Utilizing DIA-NN |  |  |  |  |  |  |  |  |
| --- | --- | --- | --- | --- | --- | --- | --- | --- |
| DIA-NN Version | Precursors Identified | Substrates Identified | CLV Substrates | HQ-CLV Substrates | NOC Substrates | HQ-NOC Substrates | Unknown Substrates | HQ- Products |
| 1.8.1 | 414,040 | 204,985 | 19,278 | 16,074 | 50,829 | 1,718 | 117,086 | 192 |
| 1.9.1 | 269,452 | 171,853 | 67,680 | 31,520 | 19,470 | 7,718 | 103,621 | 1,990 |
| 2.0 | 139,720 | 102,215 | 75,875 | 48,656 | 14,326 | 6,053 | 24,526 | 2,850 |

In the GluC substrate kinetics assay 139,720 precursors corresponding to 102,215 substrates were identified (Table 1, Table S3). Since we are assessing protease activity and endopeptidases undergo first order enzyme kinetics at low substrate concentrations,^31^ we performed secondary data analyses which mitigates noise from incorrectly assigned peaks during DIA-MS data analysis.

For each sample, we plotted the protease concentration and corresponding peptide intensity from DIA-NN outputs. We then fit each substrate in the dataset to either a horizontal straight line, which infers minimal to no cleavage of that substrate by the given protease, or a first order rate equation with a 1-parameter fit, which infers the disappearance of a substrate due to protease cleavage by the protease (Equation 1)

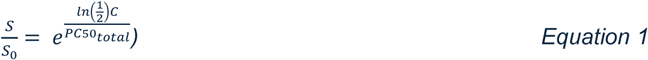

where *S* is the processed signal count of a single substrate, *S*_0_ is the median processed signal count of the substrate without any protease added, *C* is the protease concentration (nM), and *PC*_50 *total*_ is the protease concentration at which the substrate signal count is at 50% (S/S_0_ = 0.5). The horizontal straight line is normalized to *S*_0_ as well. For the first order rate equation, we are fitting for *PC*50_*total*_. The two possible fits provide statistical metrics which were then used to determine if a substrate is cleaved or not cleaved by the protease. Once substrates were designated as either cleaved or not cleaved, a stringent statistical metric criterion was used to identify a sub selection that were deemed ‘High Quality’. In the GluC substrate kinetic experiment there were 48,656 High Quality Cleaved (HQ-CLV) substrates and 6,053 High Quality Not-Cleaved (HQ-NOC) substrates that fit a straight line with *p-value* < 0.005 and no missing datapoints in triplicate LCMS acquisitions at each protease concentration. (Figure 2B, Table 1).

The next step of the data processing pipeline exploits the ability to re-analyze DIA-MS acquisitions with a different DIA assay library to look for the appearance of C-terminal peptide products after proteolytic cleavage. First, all possible C-terminal cleavage products from the HQ-CLV substrates results were computationally generated, appended to the FASTA file, and DIA-NN was re-run in library free mode looking for C-terminal cleavage products generated in the protease cleavage experiment. Secondary data analysis was performed fitting all observed C-terminal cleavage products to a 2-parameter fit of the inverse of Eqn 1.

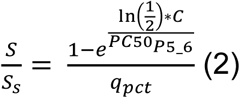

Where *S*_*s*_ is the median processed signal intensity at the highest protease concentration, *PC*50_*P*5_6_ is the protease concentration at which 50% of the maximum processed signal was observed, and *q*_*pct*_ is the cleavage percentage parameter. In many cases, cleaved products do not reach the saturated signal limit at the highest protease concentration, which does not represent the characteristics of a sigmoidal curve. To account for this, we added the parameter, *q*_*pct*_, to fit for the theoretical plateau (e.g., saturated signal level) for a given cleaved product. We constrained the fitting of *q*_*pct*_ between 0.5 and 1.4 to ensure the fitting does not deviate to extreme ranges. The *PC*50_*P*5_6_ considers the cleavage rate of a specific peptide bond in the full substrate. The *PC*50_*total*_ measures the cleavage rate over all peptide bonds of the full substrate. Secondary data analysis is expanded upon in Supplementary Information.

Substrate product pairs provide valuable information when determining where a substrate is cleaved (P1 and P1’) and are helpful for aligning computational models especially when protease specificity is unknown. In the GluC substrate kinetic experiment we found 2,943 High Quality products (HQ-products) that fit Eqn 2 with a *p-value* < 0.005 and match a unique HQ-CLV substrate. This is a 1,484% increase compared to DIA-NN 1.8.1 and a 43% increase over DIA-NN 1.9.1 (Table 1).

In these 2,850 substrate product pairs we saw an enrichment of glutamic acid or aspartic acid residues in P1 (Figure 2C). This was further filtered by relative cleavage speed and the 500 substrates with the fastest PC50’s for GluC all had glutamic acid in P1 and an enrichment of hydrophobic amino acids (tyrosine, leucine, methionine, isoleucine, alanine) in P1’ (Figure 2D). Additionally, we found an excellent correlation between *PC*_50 *total*_ measurements from observed HQ-CLV substrate *PC*50_*total*_ and *PC*50_*P*5_6_ measurements from HQ-products with a slope of 0.92 and a R^2^= 0.947 (Figure 2E).

### Quantitatively Modeling Protease Specificity with a PSM

After secondary data analysis, PC50 values from HQ-CLV substrates, HQ-NOC substates, and HQ-products were used to train a PSM that quantitatively maps substrate specificity of a protease and the relative effect of amino acid substitutions at each position in a 9mer cleavage sequence. A PSM is trained as a linear model with 171 parameters for 18 possible amino acids substitutions (cysteine is excluded and alanine is the reference) in a 9mer cleavage sequence and X denoting the absence of an amino acid. In a PSM, PC50 values of a 9mer sequence are shown in log_2_ scale and alanine residues are fixed to 0. Negative values represent the predicted log_2_ PC50 reduction of an amino acid substitution relative to alanine, i.e. how much less protease is needed to cleave the alanine substituted peptide. Positive values represent an amino acid substitution predicted to require higher protease concentrations to cleave an alanine-substituted substrate.

PSM models were trained using 80% of the data. We observed that the models provide good predictions of PC50 values for both the training data and the 20% testing data (Supplementary Information, Figure S6). This suggests PSM models can be used to predict protease cleavage kinetics for any 9mer peptide sequence.

Generating a PSM for the GluC substrate kinetics experiment (Figure 2F), we saw a strong preference for glutamic acid at P1, with a −12.75 log_2_ PC50 (∼6,888 fold) increase in cleavability of a substrate compared to alanine. Aspartic acid has a −4.02 log_2_ PC50 (∼16 fold) increase in cleavability of a substrate versus alanine corroborating published data^17,18^ of GluC’s ∼400x preference of glutamic acid over aspartic acid at P1 (Figure 2F). According to the PSM the fastest predicted GluC substrate is “WWVLE-MFFL” predicted to be 6,813,667x faster than a 9mer sequence of alanine. Interestingly, a GluC sequence with a glutamic acid at P1 but a proline in P1’ is predicted to be cleaved 3,848x slower than an alanine which can be further slowed down another 467x with the addition of proline in P2’, negating any benefit of glutamic acid at P1. Proline in P3 confers a ∼25x decrease in cleavability however a proline in P2 or P1 has marginal effect on GluC’s proteolytic activity. We also observe a 24x and 69x reduction in cleavability for “E-E” and “E-D” bonds respectively showing two negatively charged residues at P1 and P1’ hinders GluC’s function.

A PSM can predict the speed of any 9 amino acid cleavage sequence including those not measured in during library analysis. To test the accuracy of PC50 predictions, we calculated PC50’s for all substrates measured in the assay. (Figure 2G) The predictions correlate well with experimental data (slope=0.91, R^2^=0.901) and the linear regression fit difference between measured and predicted PC50 values were negligible between training and testing data for GluC (Figure S6A-B) indicating the linear model-based PSM accurately maps GluC’s substrate specificity.

### Mapping the Dual Function Protease Fibroblast Activated Protein

FAP is a complex protease known to function both as an endopeptidase and dipeptidyl exopeptidase. ^19,21,32,33^ Similarly to GluC, we performed a FAP substrate kinetic assay with the naive substrate library. Triplicate diaPASEF injections for each protease concentration were acquired and data was analyzed as described for GluC. Secondary data analysis curve fittings resulted in the identification of 15,569 HQ-CLV substrates, 12,576 HQ-NOC substrates, and 4,948 HQ-products. (Figure 3A, Table S4) Of these, 1,203 substrate product pairs showed endopeptidase activity with an enrichment of glycine in P2 and proline or alanine in P1 (Figure 3B). We observe subtle differences in the 500 fastest substrates with endopeptidase activity with a greater enrichment of glycine in P2. 3,422 substrates showed dipeptidyl peptidase activity and an enrichment of glycine in P2 and alanine and serine in P1(Figure 3D). The 500 fastest substrates with dipeptidyl activity show a similar enrichment of glycine in P2 but show a stronger preference for alanine in P1(Figure 3E).

**Figure 3:**
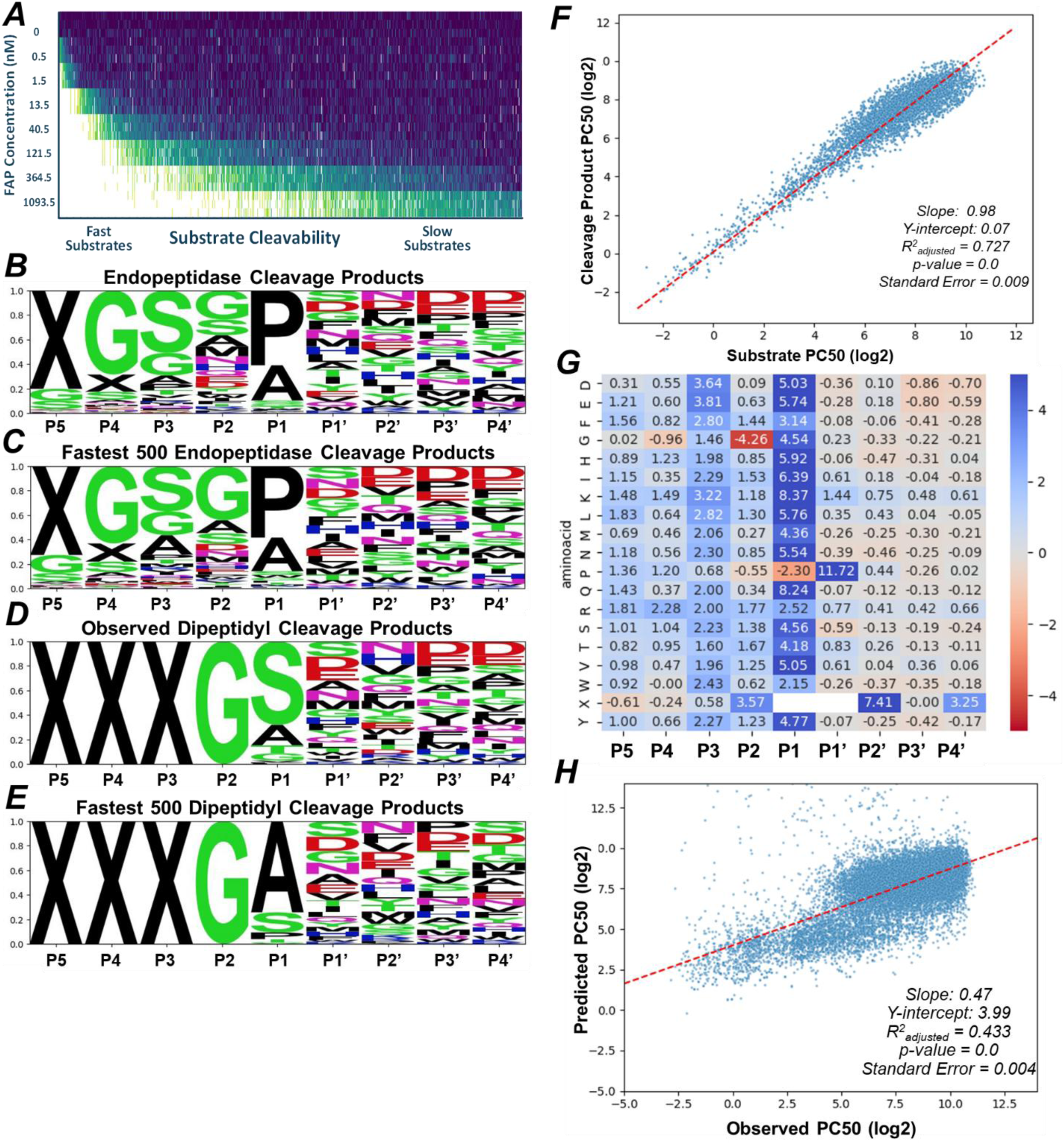
**(A)** Heatmap of high quality cleaved substrates in FAP experiment showing a gradation of very fast substrates to relatively slow substrates cleaved by FAP **(B)** SeqLogo of 1,526 substrate product pairs cleaved by FAP as an endopeptidase **(C)** SeqLogo of the 500 fastest substrate product pairs cleaved by FAP as an endopeptidase **(D)** SeqLogo of 3,422 substrate product pairs cleaved by FAP as an dipeptidyl exopeptidase **(E)** SeqLogo of the 500 fastest substrate product pairs cleaved by FAP as an dipeptidyl exopeptidase **(F)** Scatterplot of 4,948 substrates where we could detect high quality products (Y Axis) and high quality cleaved substrates (X Axis) showing the correlation of PC50 measurements of each. Pairs with absolute difference > 2 were considered outliers and removed from analysis **(H)** Positional Scanning Matrix (PSM) of FAP experiment characterizing FAP’s substrate specificity. All amino acids are fixed to Alanine; a negative number represents the log2 PC50 speed an amino acid substitution will speed up a substrate and a positive number will slow down the substrate in respect to Alanine. X is indicative of a lack of a residue **(G)** Scatterplot showing PSM predicted PC50’s (Y-Axis) vs Observed PC50’s (X-Axis) for the FAP substrate kinetics experiment with the naïve substrate library. Predicted PC50’s correlated with observed PC50’s with a R2=0.901 and slope = 0.91 **(H)** Scatterplot showing PSM predicted PC50’s (Y-Axis) vs Observed PC50’s (X-Axis) for the FAP substrate kinetics experiment.

We found a good correlation between *PC*_50 *total*_ measurements from observed HQ-CLV substrate *PC*50_*total*_ and *PC*50_*P*5_6_ measurements from HQ-products (slope=0.98, R^2^=0.727, Figure 3F). A PSM was then generated showing the enrichment of P2 glycine and P1 proline. Minimal effects were seen in the P1’-P4’ residues aside from a P1’ proline resulting in a 3,373x disruption in FAP’s proteolytic activity (Figure 3G). FAP’s PSM predicts the fastest cleavage sequence is “XGAGP-SHDD” which is ∼55x faster than the previously reported fastest FAP substrate, “ASGP-SS”, obtained from targeted peptide synthesis experiments.^19^ PC50’s were next predicted for all substrates measured in the assay (Figure 3H). The predictions correlate with experimental data (slope=0.47, R^2^=0.433) and the linear regression fit difference between measured and predicted PC50 values were negligible between training and testing data for FAP (Figure S6C-D) indicating the linear model-based PSM is sufficient to map substrate specificity from a dual function protease like FAP.

## Discussion

We have developed a highly multiplexed screen that can quantitatively map protease substrate specificity in a single unbiased assay. This LC/MS based naïve peptide substrate library approach results in substrate kinetic rates for thousands of substrates per protease and a PSM that can be used to predict the cleavage rate for any sequence, including those not assayed in the library.

We applied this method to GluC, a well-studied protease with defined substrate specificity and generated a PSM sufficient to quantitatively map GluC’s substrate specificity. Although GluC is useful in proteomics style experiments and in multi-enzyme peptide mapping of therapeutic antibodies,^17,34^ all peptides containing a glutamic acid are not cleaved equally by GluC, especially in situations where glutamic acid is followed by one or more prolines. When GluC was characterized, it was noted a proline P1’ significantly reduces cleavage.^35^ In the MEROPS database there are 4,330 cleavage sequences cataloged for GluC and only 8 have a proline in P1’. With the data generated here we can quantitatively map the detrimental effect of a proline in P1’, P2’, and P3 as well as the inhibitory effect of two negatively charged residues in both P1 and P1’ on GluC’s activity expanding upon what is known about this well-studied protease in a single unbiased assay. Integrating HQ-NOC substates into a PSM provides significant value especially in scenarios where proline or two negatively charged residues block GluC’s proteolytic activity. Using the GluC PSM, one could generate in-silico substrates predicted to be cleaved by low concentrations of GluC and use these fast GluC peptides as a sensor when assessing a biologic during lead optimization or any purpose requiring GluC releasable peptides.

Subsequently, we applied this method to FAP, a complex protease functioning as both a dipeptidyl exopeptidase and endopeptidase. In a single unbiased assay, we were able to quantitatively map FAP’s dual function substrate specificity. If one were to assay FAP’s substrate specificity using phage or RNA display, this information would be lost. The naïve substrate library was not constructed to assay dipeptidyl exopeptidases as it has a fixed N-terminal glycine but is still powerful enough to generate quantitative kinetic curves of thousands of FAP cleavable substrates in its function as a dipeptidyl and endopeptidase adding substantially to the 10 FAP substrates currently available in MEROPS.^20^

Utilizing a mass spectrometer for this assay has significant benefits over phage- and RNA-display approaches ^7,13–15^ and current LC/MS based methods ^11,12,369,11,12^ with few downsides. The method described here is faster and more easily scalable than semi-quantitative phage- and RNA-display approaches that result in enriched substrate motifs and rely on an additional peptide synthesis step and LC/MS measurement for each protease assayed to determine substrate kinetics. When comparing the naïve peptide substrate library approach to proteome-derived peptide libraries we see significant gains in mapping protease substrate specificity. Assaying GluC using a proteome-derived peptide library, other than an enrichment of glutamic acid and aspartic acid in P1, no selectivity for any other residue between P6–P6′ was observed^12^ whereas the GluC PSM generated here can quantitatively interrogate subtle effects of amino acid substitutions at each position in a cleavage sequence. Unlike proteome-based libraries which rely on peptides found in nature, the naïve substrate library described here was deliberately designed to elute over the LC gradient, have an even ion mobility distribution, and unambiguously differentiate each peptide in the library. Additionally, proteome-derived peptide libraries have a dynamic range of ∼7 orders of magnitude.^16^ Despite minor variations in cloning efficiency and competitive grown advantages during *E. coli* production, the narrow dynamic range in the substrate library allows for 200,000 substrates to be multiplexed, providing sufficient peptide diversity to quantitatively map protease specificity in a single assay.

To our knowledge this is the first time a peptide library has been generated at this scale from a carrier protein produced in *E. coli*, an organism absent from most PTM’s including glycosylation which can affect protease activity.^37,38^ We used a XTENylated Bi-specific T-cell engager carrier protein. The very high solubility of XTEN facilitates the expression of fusion sequences with low solubility. ^39^ In addition, XTEN allows efficient purification via ion exchange. This robust approach resulted in a naïve substrate library capable of quantitatively mapping substrate specificities of multiple proteases utilizing the same set of peptides, having the potential to identify subtle differences in substrate specificity between related proteases.

There are, however, some downsides to our approach. Although the library of ∼200,000 substrates can be analyzed in a single experiment, pushing LC/MS technology to its limits, this approach cannot assess the complete search space for a 9mer cleavage sequence that can be achieved in RNA-display.^15^ Nevertheless, PSM’s generated by this approach are sufficiently powered to predict cleavage speed of substrates for GluC and FAP. Assaying the complete search space may not be necessary to map protease specificity, especially when coupled with highly quantitative multiplexed substrate kinetics as has been done here.

In conclusion, the method described here provides the framework to determine substrate specificity of a protease in a single LC/MS experiment. A naïve peptide substrate library was designed, produced in *E. coli*, and assayed with both GluC and FAP resulting in thousands of substrate kinetics curves for each protease. Quantitative substrate specificity maps were generated for both proteases robust enough to accurately predict the relative cleavage rate of peptide sequences for their respective protease, significantly expanding our knowledge about substrate specificity for both proteases. The naïve substrate library approach described here should be able to quantitatively map the specificity of and generate PSM’s for most proteases.

## Methods

### Generation, Purification, and Production of Peptide Substrate Library

Library sequences flanked by known canonical Thrombin and Factor Xa helper protease sites (LVPR-GS and IEGR-G, respectively) were cloned as a pool into the unstructured XTEN polypeptide region to the N terminus of an EGFR T-Cell Engager,^26^ used only as a carrier protein during purification and not for its function as a T-Cell Engager. The mixture was produced as previously described.^28^ Briefly the mixture was fermented in *E. coli* and prepared as a cell pellet. The cell pellet was resuspended until homogenous, mechanically lysed, flocculated, then submicron filtered. The supernatant was processed through a series of anion and cation chromatography resins, and filtration. The purified protein mixtures containing the naïve substrate library regions were then digested into a peptide library using either Thrombin or Factor Xa. The resulting mixture underwent chromatography clean-up using anion exchange, followed by benzamidine affinity chromatography targeting serine proteases to remove residual Thrombin or Factor Xa. The resulting naïve substrate library was desalted and concentrated using a HLB sorbent (Waters) and used for subsequent experiments to map protease specificity.

### GluC Substrate Kinetics Assay

Naïve substrate library screened with GluC (Roche 11420399001) was performed as follows: (1) Started with 1093nM concentration of GluC in reaction buffer,10mM CaCl2, 50mM Tris, pH 7.5, and made 8 3-fold dilutions, as well as a no protease control sample. (2) Spike in naïve substrate library to each sample and incubate at 37°C shaking for 24 hours. (3) Acidify each sample with formic acid to stop protease activity. (4) Inject samples in triplicates onto Vanquish Neo UHPLC system (Thermo) coupled to a timsTOF HT mass spectrometer (Bruker) and acquire diaPASEF data.

### FAP Substrate Kinetics Assay

Naïve substrate library screened with FAP (Abcam ab79623) was performed as follows: (1) Started with 1093nM concentration of FAP in reaction buffer,10mM MgCl2, 1mM MnCl2, 50mM Tris, pH 7.5, and made 8 3-fold dilutions, as well as a no protease control sample. (2) Spike in naïve substrate library to each sample and incubate at 37°C shaking for 24 hours. (3) Acidify each sample with formic acid to stop protease activity. (4) Inject samples in triplicates onto Vanquish Neo UHPLC system (Thermo) coupled to a timsTOF HT mass spectrometer (Bruker) and acquire diaPASEF data.

### LC/MS Methods

MS acquisition: MS data were acquired on timsTOF HT (Bruker) mass spectrometer operating in diaPASEF mode. The timsTOF HT mass spectrometer was equipped with the CaptiveSpray ion source and connected to the Vanquish Neo UHPLC system (Thermo Fisher Scientific) operating at a flow rate of 1 µL/min using mobile phase Solvents A (0.1% formic acid in water) and B (0.1% formic acid in acetonitrile), an inline PepMap Neo C18 trap cartridge (5 mm x 0.3 mm x 5 µm particle size, Thermo Fisher Scientific), a nanoEase M/Z Peptide BEH C18 analytic column (15 cm x 300 µm x 1.7 µm particle size, Waters) housed in the Vanquish column compartment at 50°C, and a 20 µm Classic Emitter (Bruker) with a linear gradient of 3-6%B for 2 minutes, 6-30%B for 80 minutes, 30-50%B for 2 minutes, 50-95%B for 0.5 minutes, holding at 95% and re-equilibrating to 3%B for a 100 min total. For diaPASEF acquisition on the timsTOF HT, the MS scan range was 100–1700 m/z, the ion mobility window was 1/K0 0.7–1.23, the ramp and accumulation time was 100 ms, the source voltage was 1700 V, the dry gas was 3.0-L/min, and the dry temperature was 200 ◦C. The mobilogram shape and ion mobility windows were tailored to the naïve substrate library.

The mass spectrometry proteomics data and DIA-NN outputs have been deposited to the ProteomeXchange Consortium via the PRIDE^40^ partner repository with the dataset identifier PXD061955

### DIA-NN Methods

Raw data processing was carried out with DIA–NN v.1.8.1, DIA–NN v.1.9.1, and DIA– NN v.2.0. For DIA-NN 1.8.1 default settings in ‘robust LC (high accuracy)’ and ‘--cut ‘ was used. For DIA–NN v.1.9.1 ‘QuantUMS High Precision’ and ‘--cut ‘ were enable. For DIA–NN v.2.0 ‘QuantUMS High Precision’, --proteoforms, ‘--cut ‘, and ‘--dg-keep-cterm 4’ were enabled.

## Acknowledgements

The authors would like to thank Ardigen for their help in building a database to store and analyze these complex datasets.

## References

1. Puente, X. S., Sánchez, L. M., Overall, C. M. & López-Otín, C. Human and mouse proteases: a comparative genomic approach. Nat. Rev. Genet. 4, 544–558 (2003).

2. Urban, S. Making the cut: central roles of intramembrane proteolysis in pathogenic microorganisms. Nat. Rev. Microbiol. 7, 411–423 (2009).

3. Reed, C. E. & Kita, H. The role of protease activation of inflammation in allergic respiratory diseases. J. Allergy Clin. Immunol. 114, 997–1008 (2004).

4. Mason, S. D. & Joyce, J. A. Proteolytic networks in cancer. Trends Cell Biol. 21, 228–237 (2011).

5. Nagasawa, T., Kawaguchi, M., Nishi, K. & Yasumasu, S. Molecular evolution of hatching enzymes and their paralogous genes in vertebrates. BMC Ecol. Evol. 22, 9 (2022).

6. Eckhard, U. et al. Active site specificity profiling of the matrix metalloproteinase family: Proteomic identification of 4300 cleavage sites by nine MMPs explored with structural and synthetic peptide cleavage analyses. Matrix Biol. 49, 37–60 (2016).

7. Zhou, J. et al. Deep profiling of protease substrate specificity enabled by dual random and scanned human proteome substrate phage libraries. Proc. Natl. Acad. Sci. 117, 25464–25475 (2020).

8. Cieplak, P. & Strongin, A. Y. Matrix metalloproteinases – From the cleavage data to the prediction tools and beyond. Biochim. Biophys. Acta (BBA) - Mol. Cell Res. 1864, 1952–1963 (2017).

9. Backes, B. J., Harris, J. L., Leonetti, F., Craik, C. S. & Ellman, J. A. Synthesis of positional-scanning libraries of fluorogenic peptide substrates to define the extended substrate specificity of plasmin and thrombin. Nat. Biotechnol. 18, 187–193 (2000).

10. Rohweder, P. J., Jiang, Z., Hurysz, B. M., O’Donoghue, A. J. & Craik, C. S. Multiplex substrate profiling by mass spectrometry for proteases. Methods Enzym. 682, 375–411 (2022).

11. O’Donoghue, A. J. et al. Global identification of peptidase specificity by multiplex substrate profiling. Nat. Methods 9, 1095–1100 (2012).

12. Schilling, O. & Overall, C. M. Proteome-derived, database-searchable peptide libraries for identifying protease cleavage sites. Nat. Biotechnol. 26, 685–694 (2008).

13. Kretz, C. A., Tomberg, K., Esbroeck, A. V., Yee, A. & Ginsburg, D. High throughput protease profiling comprehensively defines active site specificity for thrombin and ADAMTS13. Sci. Rep. 8, 2788 (2018).

14. Ratnikov, B. I., Cieplak, P., Remacle, A. G., Nguyen, E. & Smith, J. W. Quantitative profiling of protease specificity. PLoS Comput. Biol. 17, e1008101 (2021).

15. Zhu, B. et al. Comprehensive protease specificity profiling. bioRxiv 2024.11.06.622033 (2024) doi:10.1101/2024.11.06.622033.

16. Zubarev, R. A. The challenge of the proteome dynamic range and its implications for in-depth proteomics. PROTEOMICS 13, 723–726 (2013).

17. Swaney, D. L., Wenger, C. D. & Coon, J. J. Value of Using Multiple Proteases for Large-Scale Mass Spectrometry-Based Proteomics. J. Proteome Res. 9, 1323–1329 (2010).

18. Breddam, K. & Meldal, M. Substrate preferences of glutamic-acid-specific endopeptidases assessed by synthetic Peptide Substrates based on intramolecular fluorescence quenching. Eur. J. Biochem. 206, 103–107 (1992).

19. Edosada, C. Y. et al. Peptide substrate profiling defines fibroblast activation protein as an endopeptidase of strict Gly2-Pro1-cleaving specificity. FEBS Lett. 580, 1581–1586 (2006).

20. Rawlings, N. D., Barrett, A. J. & Bateman, A. MEROPS: the peptidase database. Nucleic Acids Res. 38, D227–D233 (2010).

21. Keane, F. M., Nadvi, N. A., Yao, T. & Gorrell, M. D. Neuropeptide Y, B-type natriuretic peptide, substance P and peptide YY are novel substrates of fibroblast activation protein-α. FEBS J. 278, 1316–1332 (2011).

22. Laronha, H. & Caldeira, J. Structure and Function of Human Matrix Metalloproteinases. Cells 9, 1076 (2020).

23. Yadati, T., Houben, T., Bitorina, A. & Shiri-Sverdlov, R. The Ins and Outs of Cathepsins: Physiological Function and Role in Disease Management. Cells 9, 1679 (2020).

24. Meier, F. et al. diaPASEF: parallel accumulation–serial fragmentation combined with data-independent acquisition. Nat. Methods 17, 1229–1236 (2020).

25. Gessulat, S. et al. Prosit: proteome-wide prediction of peptide tandem mass spectra by deep learning. Nat. Methods 16, 509–518 (2019).

26. Plante, P.-L. et al. Predicting Ion Mobility Collision Cross-Sections Using a Deep Neural Network: DeepCCS. Anal. Chem. 91, 5191–5199 (2019).

27. Escher, C. et al. Using iRT, a normalized retention time for more targeted measurement of peptides. PROTEOMICS 12, 1111–1121 (2012).

28. Cattaruzza, F. et al. Precision-activated T-cell engagers targeting HER2 or EGFR and CD3 mitigate on-target, off-tumor toxicity for immunotherapy in solid tumors. Nat. Cancer 4, 485–501 (2023).

29. Demichev, V., Messner, C. B., Vernardis, S. I., Lilley, K. S. & Ralser, M. DIA-NN: neural networks and interference correction enable deep proteome coverage in high throughput. Nat. Methods 17, 41–44 (2020).

30. Kistner, F., Grossmann, J. L., Sinn, L. R. & Demichev, V. QuantUMS: uncertainty minimisation enables confident quantification in proteomics. bioRxiv 2023.06.20.545604 (2023) doi:10.1101/2023.06.20.545604.

31. Duggleby, R. G. [3] Analysis of enzyme progress curves by nonlinear regression. Methods Enzym. 249, 61–90 (1995).

32. Hamson, E. J., Keane, F. M., Tholen, S., Schilling, O. & Gorrell, M. D. Understanding fibroblast activation protein (FAP): Substrates, activities, expression and targeting for cancer therapy. Proteom. Clin. Appl. 8, 454–463 (2014).

33. Edosada, C. Y. et al. Selective Inhibition of Fibroblast Activation Protein Protease Based on Dipeptide Substrate Specificity*. J. Biol. Chem. 281, 7437–7444 (2006).

34. Pradhan, G. et al. Multiple-parallel-protease digestion coupled with high-resolution mass spectrometry: An approach towards comprehensive peptide mapping of therapeutic mAbs. J. Proteom. 232, 104053 (2021).

35. Drapeau, G. R. [38] Protease from Staphyloccus aureus. Methods Enzym. 45, 469–475 (1976).

36. Vizovišek, M., et al. Protease Specificity: Towards In Vivo Imaging Applications and Biomarker Discovery. Trends Biochem. Sci. 43, 829–844 (2018).

37. Madzharova, E., Sabino, F., Kalogeropoulos, K., Francavilla, C. & Keller, U. auf dem. Substrate O-glycosylation actively regulates extracellular proteolysis. Protein Sci. 33, e5128 (2024).

38. Walsh, G. & Jefferis, R. Post-translational modifications in the context of therapeutic proteins. Nat. Biotechnol. 24, 1241–1252 (2006).

39. Schellenberger, V. et al. A recombinant polypeptide extends the in vivo half-life of peptides and proteins in a tunable manner. Nat. Biotechnol. 27, 1186–1190 (2009).

40. Perez-Riverol, Y. et al. The PRIDE database resources in 2022: a hub for mass spectrometry-based proteomics evidences. Nucleic Acids Res. 50, D543–D552 (2021).

41. Meier-Abt, F. et al. The protein landscape of chronic lymphocytic leukemia. Blood 138, 2514–2525 (2021).

42. Demichev, V. et al. dia-PASEF data analysis using FragPipe and DIA-NN for deep proteomics of low sample amounts. Nat. Commun. 13, 3944 (2022).

43. Chang, J. Thrombin specificity. Eur. J. Biochem. 151, 217–224 (1985).

44. Brandstetter, H. et al. X-ray Structure of Active Site-inhibited Clotting Factor Xa IMPLICATIONS FOR DRUG DESIGN AND SUBSTRATE RECOGNITION*. J. Biol. Chem. 271, 29988–29992 (1996).

