## Supplemental Information for "Quantitative Substrate Kinetics Screening: One-Pot LC/MS Based Approach to Map Protease Specificity"

### Supplementary Information

#### Optimizing Data Quality Utilizing DIA-NN

Recent improvements in the DIA-NN software package, including most recently QuantUMS[30] in DIA-NN 1.9.1 and custom decoys implemented in DIA-NN 2.0 for datasets with fixed N- or C- terminal amino acids, have resulted in significant improvements in data quality, magnified further with the complex data processing pipeline we have established here. QuantUMS integrates multiple signals associated with each precursor, and custom decoy models has significant benefit as the naïve substrate library contains fixed -LVPR and -GIEGR c-terminal amino acids.

When comparing DIA-NN 1.8.1, 1.9.1, and 2.0, we see considerable differences in total precursor and substrate identifications. DIA-NN 1.8.1 identifies 414,040 precursors and 204,985 substrates, DIA-NN 1.9.1 identifies 269,452 precursors and 171,853 substrates, and DIA-NN 2.0 identifies 139,720 precursors and 102,215 substrates. Strikingly, after secondary data analysis, we see the opposite trend with DIA-NN 2.0 identifying 48,656 High Quality Cleaved Products a 1.5x increase in high quality data when compared to DIA-NN 1.9.1 and a 3x increase when compared to DIA-NN 1.8.1.(Table 1) We see a 4.5x increase in the number of High Quality Not-Cleaved Substrates between DIA-NN 1.8.1 and DIA-NN 1.9.1 which isn't significantly changed in DIA-NN 2.0. Additionally, the number of Unknown substrates, which don't fit either the first order rate equation or a straight line, decrease substantially between DIA-NN 1.8.1, DIA-NN 1.9.1, and DIA-NN 2.0 and due to enzyme kinetics can be considered noise or misassigned peaks in our dataset.

When comparing the PC50 distribution of all the High-Quality Cleaved Substrates between the same GluC substrate kinetics assay dataset analyzed with the different DIA-NN versions, there is a population of substrates that are cleaved at low protease concentrations that was missed with both DIA-NN 1.8.1 and DIA-NN 1.9.1. (Figure S5A-C) Although the total number of precursors identified in the experiment has decreased, there are significantly more of these substrates which fulfill the stringent filtering criteria after secondary data analysis when analyzed with QuantUMS and the Decoy Strategy introduced in DIA-NN 2.0 (Table 1) and there is less data missingness and incorrectly assigned peaks (Figure S5D-E). Therefore, we use DIA-NN 2.0 for primary analysis of this complex naive substrate library data.

#### Secondary Data Analysis (PC50 Calculation and Classification and PSM)

Each cleavage rate experiment went through a 5-step computational pipeline to determine high-quality substrate and corresponding products. 1) **Processed samples**

**with DIA-NN to identify substrates.** The raw DIA mass spectrometry data for all the samples (.d format) were processed through DIA-NN 2.0 to identify all possible substrates and their corresponding signal. The input includes all raw data .d sample files and the FASTA file containing all full-length substrates in the combined naïve substrate sub-libraries. For this processing, we set the q-value at 0.01, missed cleavages at 1, mass accuracy at 15, no implementation of tryptic or other enzyme cutting when producing the mass spec library file, and we did not shuffle the last 4 positions of the C-terminus for the target-decoy modeling. 2) $\boldsymbol{PC_{50\,total}}$ **Calculation.** The normalization and calculation for $PC_{50\,total}$ and $PC50_{P5\_6}$ was performed using R software. The DIA-NN output provided substrate data with corresponding MS signals for each sample. Substrates were evaluated for sufficient data points representing signal disappearance based on the first-order rate equation of protease cleavage. Quality criteria required at least 6 (of 27) data points containing signal values, at least one signal value (from triplicates) in 3 of the 9 tested protease concentrations, signal values for all triplicates at no protease concentration, and maximum MS intensity above 10% of all MS signal intensities. Qualifying substrates were normalized to the median signal of triplicate samples at no protease concentration. Each normalized substrate was fitted to either a horizontal line (representing Not-Cleaved substrate) or the first-order 1-parameter PC50 total equation (Equation 1), using the nls function with 'Gauss-Newton' solver and a starting PC50 total solution of 25. Statistical metrics were generated for both models, allowing classification of substrates into five categories: Cleaved (CLV), High-Quality Cleaved (HQ-CLV), Not Cleaved (NOC), High-Quality Not Cleaved (HQ-NOC), and Unknown (UNK). Substrates failing initial quality criteria were classified as UNK. For remaining substrates, initial CLV/NOC classification was based on comparing root mean square error (RSE) of the fits. If fitting failed for one model, the substrate was classified according to the successful model; if both fits failed, it was classified as UNK. When both fits succeeded, the model with lower RSE determined classification. Both CLV and NOC substrates required RSE values below 0.5 and p-values below 0.9. Additionally, NOC substrates needed a fraction change less than 0.4, calculated as (mean of no protease – mean of max protease concentration values)/(mean of no protease). Substrates with p-values below 0.005 in the CLV category were classified as HQ-CLV. NOC substrates were upgraded to HQ-NOC if they had p-values below 0.005, MS signals for all data points, and coefficient of variation between triplicates below 20%. **3) Generating product FASTA file.** A new FASTA file was generated for all possible products coming from HQ-CLV substrates. Each product was generated from the C-terminus of the HQ-CLV substrates ranging from 6 to 14 AA. **4)** $\boldsymbol{PC50_{P5\_6}}$ **Calculation**. The products went through a similar process as the substrates. First, they were evaluated for high data quality: at least 6 (of 27) data points containing signal values, at least one signal value (from triplicates) in 3 of the 9 test protease concentrations, maximum MS intensity above 10% of all MS signal intensities, and there must be signal values for all triplicate at the maximum protease concentration tested. If it did not meet all these criteria, then it was categorized as 'No Product'. The qualifying products were normalized to the median of the maximum protease concentration tested and then was fitted to the 2-parameter $PC50_{P5\_6}$ equation (Equation 2) using the nls function with 'Gauss-Newton' solver and a starting $PC50_{P5\_6}$ solution of 50 and cleavage parameter of 1.0. If the fit failed or if the p-value is > 0.05 or cleavage percent parameter was < 0.5 or > 1.4, the product was classified as 'No Product'. If the fit succeeded and the p-value is between 0.005 to 0.05, it was classified as 'Product'. If the p-value is less than 0.005, it was classified as 'HQ-Product'. **5) Integration of Substrate and Product PC50 data.** The last step is to merge and map all substrate and product pairs together. A cleave sequence, a 9-mer aligned where the cleavage site is between position 5 and 6, was generated for each corresponding substrate-product pairs that had a successful fit and classified as either 'CLV' substrates, 'HQ-CLV' substrates, 'Product', and 'HQ-Product'. If a substrate was mapped with multiple products, the high-quality classification takes priority. If there are multiple HQ Products mapped to a single substrate, then these are excluded for further analysis. Only $PC_{50\,total}$ values were used for training PSMs.

Each PSM model was trained on an individual cleavage-rate experiment. The model was developed in Python with using 'scipy' package for optimization. Specifically, 'scipy.optimize.minimize' is used to minimize residuals between predicted PC50 values from the PSM and measured PC50 values. During PSM training, the number of data points is equal to sum of HQ-CLV substrates, HQ-NOC substrates, and HQ-Products within an experiment. The data was split by 80% training and 20% testing. The total number of parameters in the PSM are as follows: 9 positions and 18 AA and an empty space (denoted as 'X') plus a global shift parameter (9x19 + 1 = 172 parameters). The parameters with no variance within the training data were then filtered. All 9 residues were fixed to alanine and were set to a coefficient of 0 for normalization in each position. Cysteine residues were not considered because they were not included in naïve substrate library. For HQ-CLV substrates, the training data output is the $PC_{50\,total}$. To calculate a $PC_{50\,total}$ from the PSM based on the full-length substrate, the substrate is enumerated into individual 9-mer segments. To consider cleavage sites at every possible location within the full-length substrate, we included AAAX flank on the N-terminus and XAA on the C-terminus. The $PC50_{P5_6}$was then calculated for each 9-mer segment based on the PSM, and then PC50total was then calculated by taking the inverse sum of $PC50_{P5_6}$values: 1/$PC_{50\,total}$= sum(1/ $PC50_{P5\_6}$). For HQ-NOC substrates, the training data output is denoted as 'NOC'. The PC50 prediction of $PC_{50\,total}$ from a PSM for the HQ-NOC is the same calculation as the HQ-CLV. During training, the PSM adds a penalty on the residuals if the predicted $PC_{50\,total}$ value of a HQ-NOC substrate is lower than the maximum protease concentration tested within the experiment + log2PC50 of 1.5. For HQ-Products, the training data output is the

$PC_{50\,total}$ value of the corresponding substrate, and the sequence is the 9-mer cleavage sequence of the corresponding substrate where the cleavage site is between P1’ and P1. To reduce the possibility of overfitting the model, we applied a bound for ‘X’ parameters between 0 and 8. The comparison of measured PC50 values and predicted PC50 from the PSM split between Train and Test data can be found for GluC and FAP in Figure S6A-B. Overall, we found the trained PSM can sufficiently generalize the protease motif cleavage favorability on unseen tested data.
