## Supplemental Figures for "Quantitative Substrate Kinetics Screening: One-Pot LC/MS Based Approach to Map Protease Specificity"

### Supplementary Figures

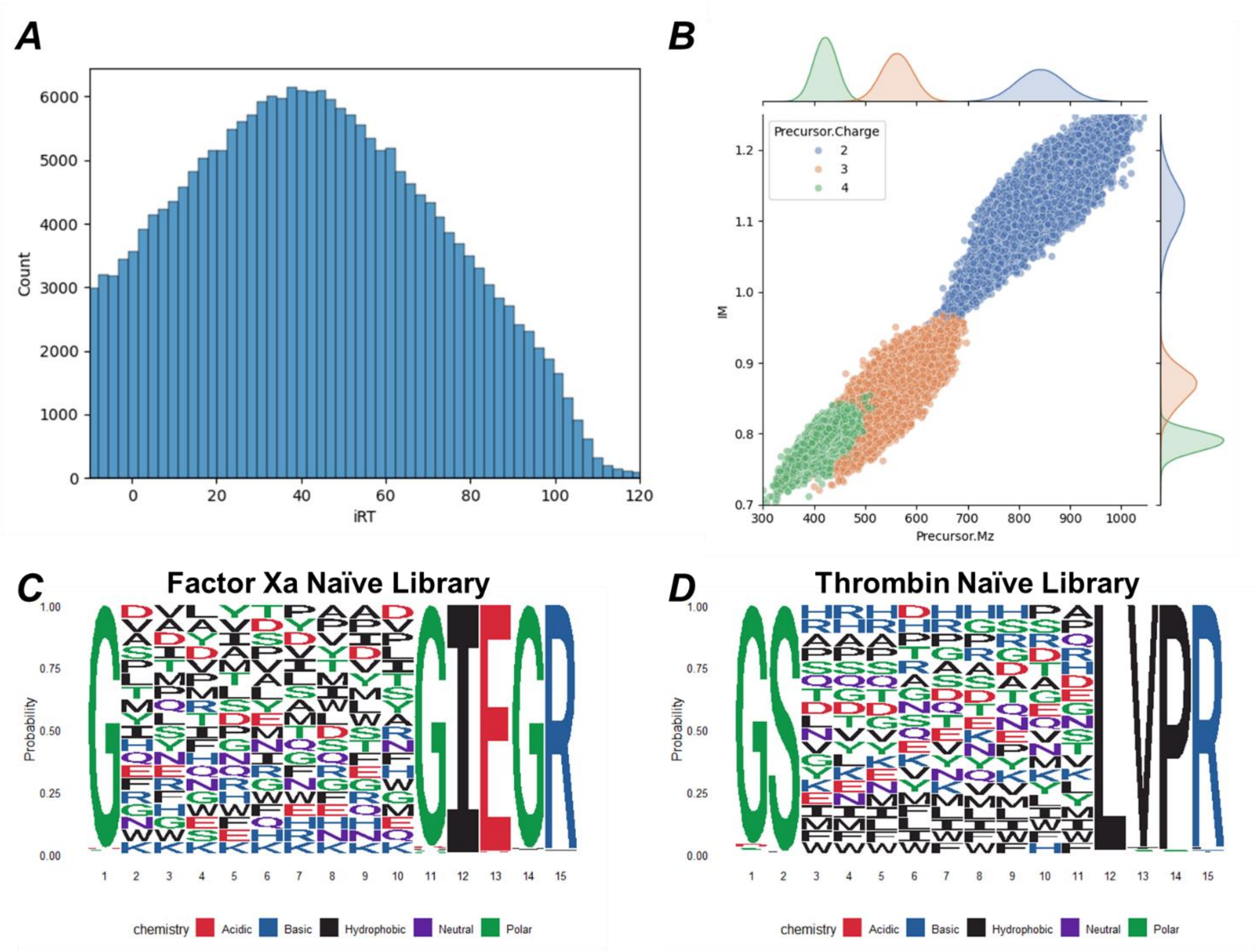


**Figure S1: (A)** Retention time prediction of all substrates in the naïve library after filtering all substrates with a IRT prediction of >-10 **(B)** Ion mobility predictions of all substrates in the naïve substrate library to ensure an even ion mobility distribution in the sample **(C)** Composition of Factor Xa naïve sub-library consisting of a fixed GIEGR-G protease cleavage remnant and a 9mer region of randomized amino acids excluding cystine **(D)** Composition of Thrombin naïve sub-library consisting of a fixed LGVP-GS protease cleavage remnant and a 9mer region of randomized amino acids excluding cystine

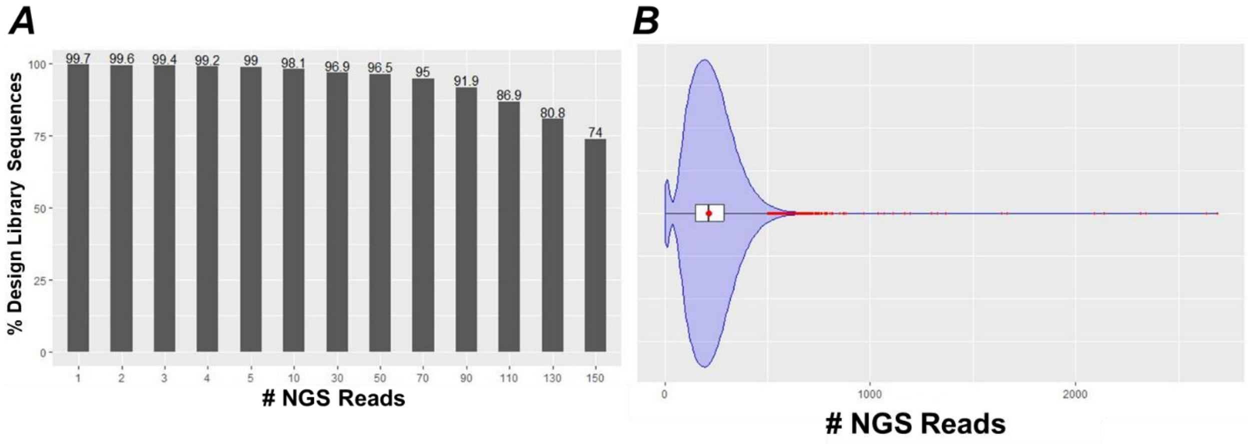


**Figure S2: (A)** Percent of designed library that was identified via NGS when looking at the number of NGS reads observed for each substrate **(B)** Violin plot of NGS reads for each substrate found in the naïve substrate library, both designed and undesigned

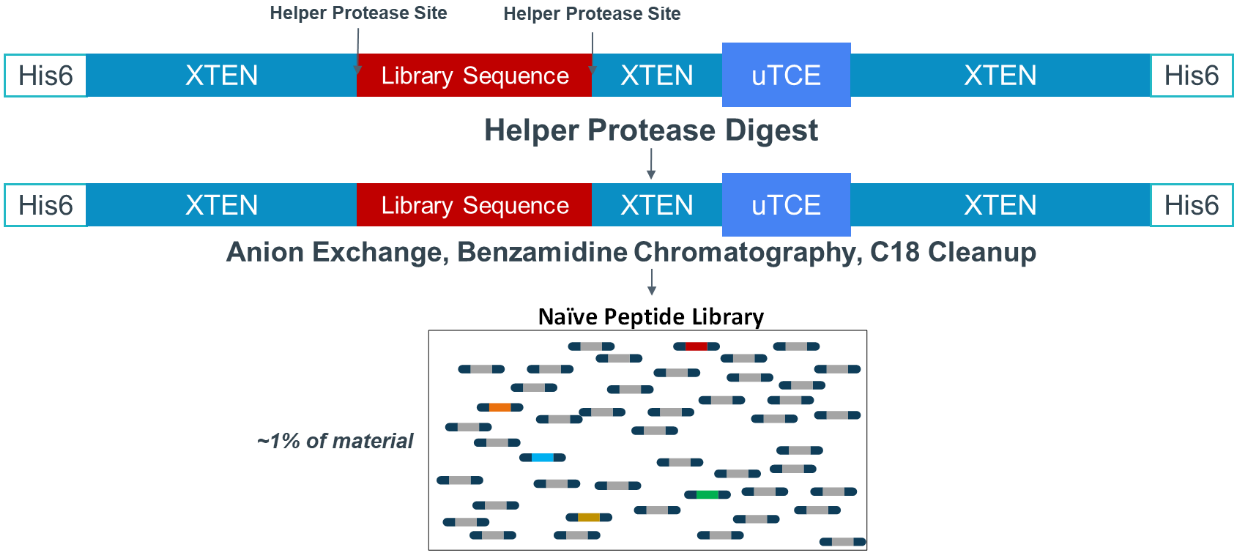


**Figure S3:** Experimental workflow to purify naïve substrate library. After E. Coli Production of a pool of XTENylated T Cell Engagers each with a unique substrate sequence, the mixture was exposed to a helper protease digest (Thrombin or Factor Xa). Anion Exchange was performed to remove free XTEN and T-Cell Engager protein. Benzamidine chromatography was then performed to remove excess helper protease and C18 cleanup was performed to remove salts and concentrate. Factor Xa and Thrombin libraries were reconstituted and pooled together.

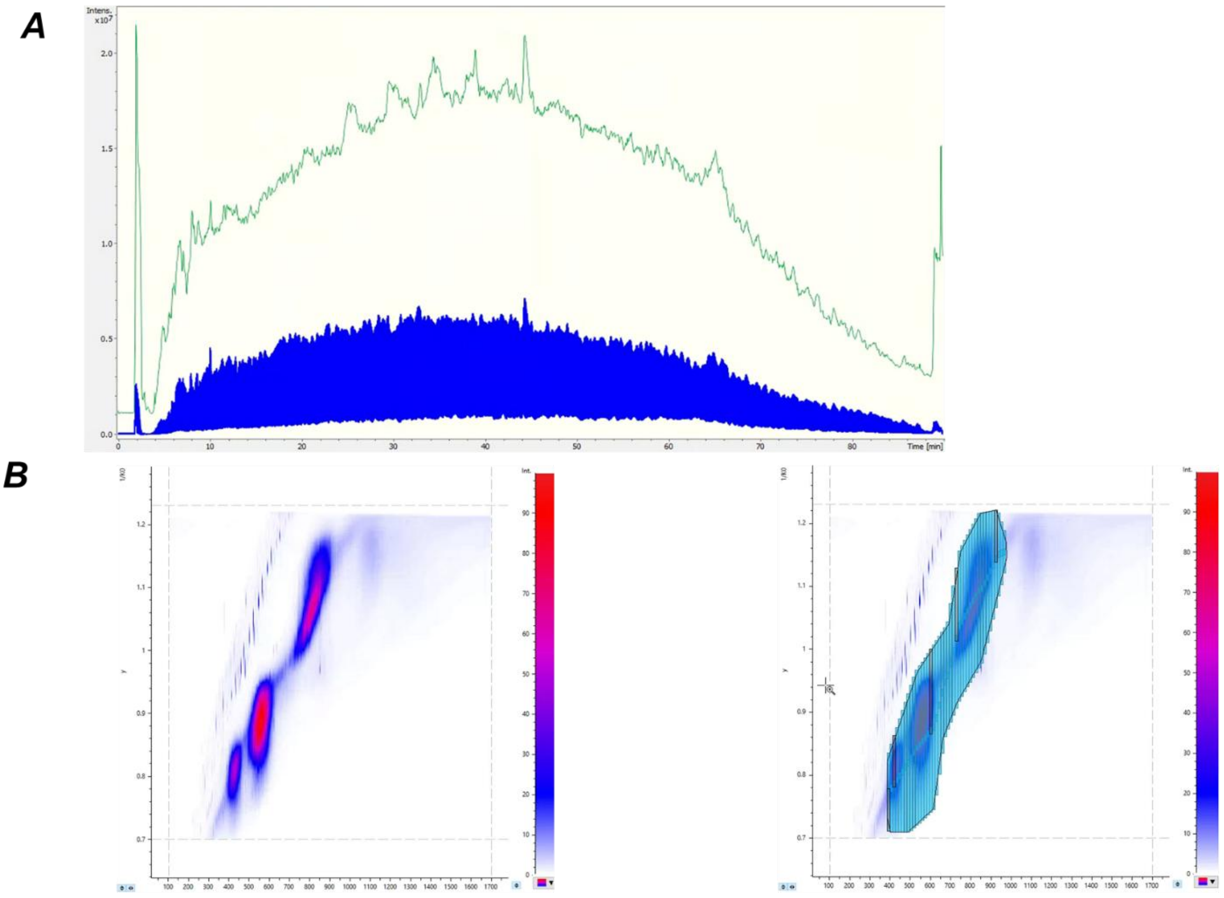


**Figure S4: (A)** TIC of a DIA-PASEF acquisition from Naïve Substrate Library without the addition of protease **(B)** Ion Mobility distribution and acquisition schema for DIA-PASEF from Naïve Substrate Library

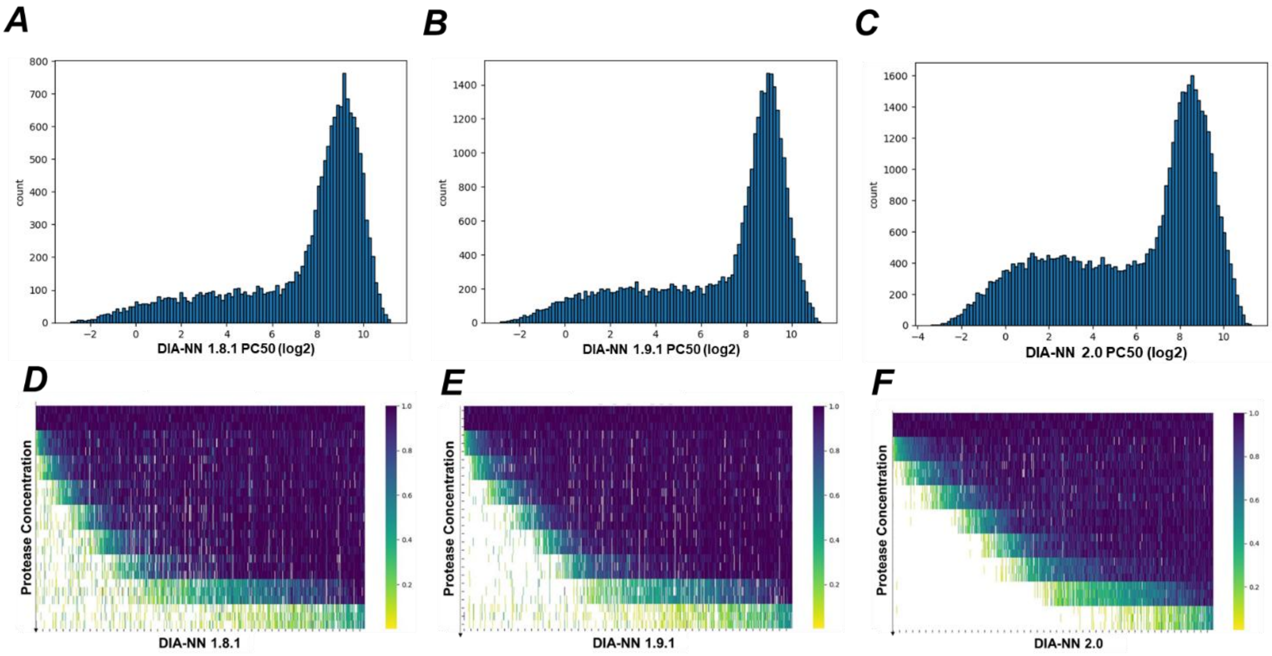


**Figure S5:** PC50 distribution of High-Quality Cleaved Substrates from the GluC substrate kinetic screen analyzed with **(A)** DIA-NN 1.8.1 **(B)** DIA-NN 1.9.1 **(C)** DIA-NN 2.0. Heatmap of the PC50 distribution of High-Quality Cleaved Substrates from the GluC substrate kinetic screen analyzed with **(D)** DIA-NN 1.8.1 **(E)** DIA-NN 1.9.1 **(F)** DIA-NN 2.0

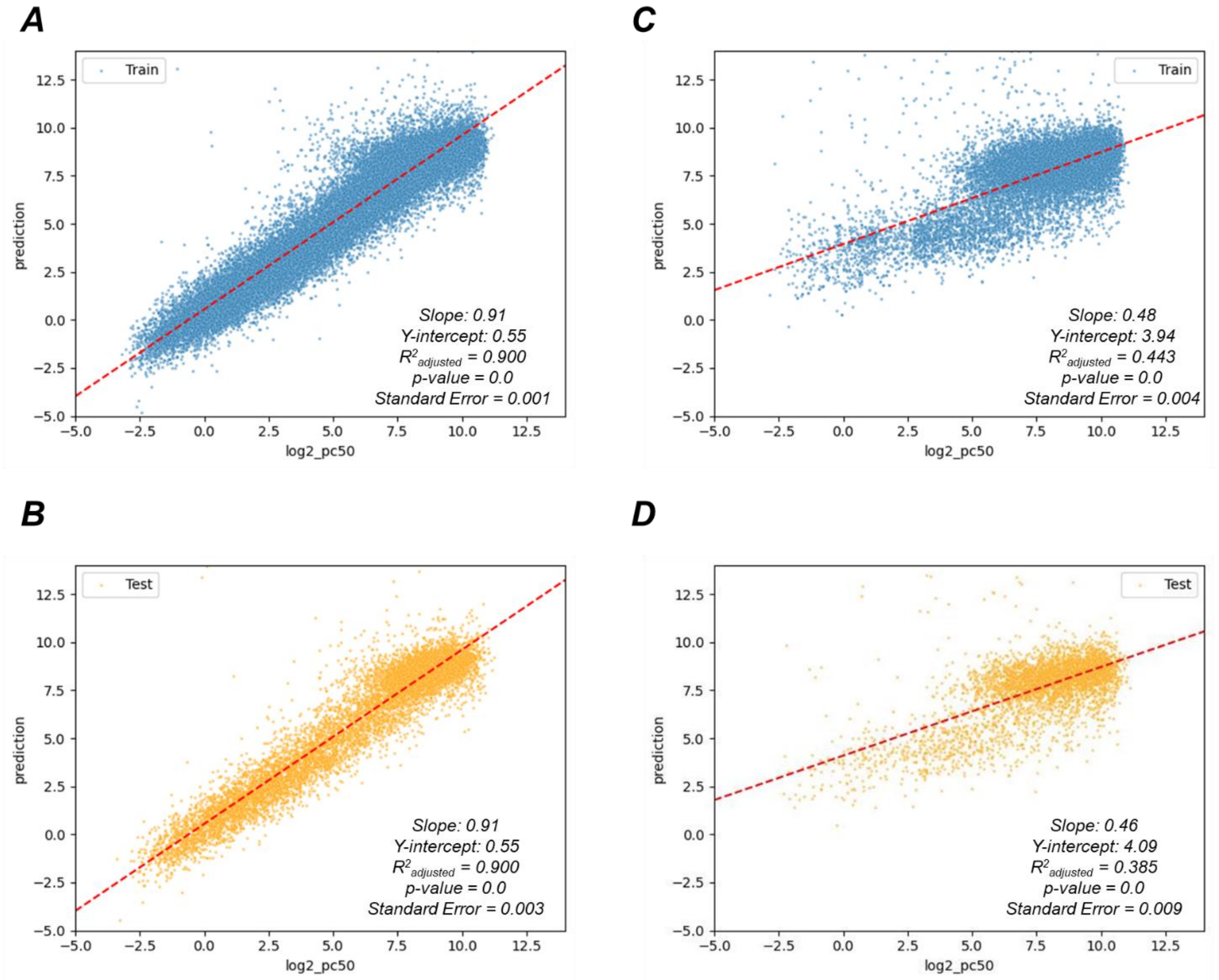


**Figure S6: (A)** Scatterplots showing PSM predicted PC50's (Y-Axis) vs Observed PC50's (X-Axis) of the training data for the GluC substrate kinetics screen with the naïve substrate library **(B)** Scatterplots showing PSM predicted PC50's (Y-Axis) vs Observed PC50's (X-Axis) of the testing data for the GluC substrate kinetics screen with the naïve substrate library **(C)** Scatterplots showing PSM predicted PC50's (Y-Axis) vs Observed PC50's (X-Axis) of the training data for the FAP substrate kinetics screen with the naïve substrate library **(D)** Scatterplots showing PSM predicted PC50's (Y-Axis) vs Observed PC50's (X-Axis) of the testing data for the FAP substrate kinetics screen with the naïve substrate library
